# Experiment-based Computational Model Predicts that IL-6 Trans-Signaling Plays a Dominant Role in IL-6 mediated signaling in Endothelial Cells

**DOI:** 10.1101/2023.02.03.526721

**Authors:** Min Song, Youli Wang, Brian H. Annex, Aleksander S. Popel

**Author notes:** Correspondence: Min Song, Department of Biomedical Engineering, School of Medicine, Johns Hopkins University, Baltimore, MD 21205.

## Abstract

Inflammatory cytokine mediated responses are important in the development of many diseases that are associated with angiogenesis. Targeting angiogenesis as a prominent strategy has shown limited effects in many contexts such as peripheral arterial disease (PAD) and cancer. One potential reason for the unsuccessful outcome is the mutual dependent role between inflammation and angiogenesis. Inflammation-based therapies primarily target inflammatory cytokines such as interleukin-6 (IL-6) in T cells, macrophages, cancer cells, muscle cells, and there is a limited understanding of how these cytokines act on endothelial cells. Thus, we focus on one of the major inflammatory cytokines, IL-6, mediated intracellular signaling in endothelial cells by developing a detailed computational model. Our model quantitatively characterized the effects of IL-6 classic and trans-signaling in activating the signal transducer and activator of transcription 3 (STAT3), phosphatidylinositol 3-kinase/protein kinase B (PI3K/Akt), and mitogen-activated protein kinase (MAPK) signaling to phosphorylate STAT3, extracellular regulated kinase (ERK) and Akt, respectively. We applied the trained and validated experiment-based computational model to characterize the dynamics of phosphorylated STAT3 (pSTAT3), Akt (pAkt), and extracellular regulated kinase (pERK) in response to IL-6 classic and/or trans-signaling. The model predicts that IL-6 classic and trans-signaling induced responses are IL-6 and soluble IL-6 receptor (sIL-6R) dose-dependent. Also, IL-6 trans-signaling induces stronger downstream signaling and plays a dominant role in the overall effects from IL-6. In addition, both IL-6 and sIL-6R levels regulate signaling strength. Moreover, our model identifies the influential species and kinetic parameters that specifically modulate the pSTAT3, pAkt, and pERK responses, which represent potential targets for inflammatory cytokine mediated signaling and angiogenesis-based therapies. Overall, the model predicts the effects of IL-6 classic and/or trans-signaling stimulation quantitatively and provides a framework for analyzing and integrating experimental data. More broadly, this model can be utilized to identify targets that influence inflammatory cytokine mediated signaling in endothelial cells and to study the effects of angiogenesis- and inflammation-based therapies.

## INTRODUCTION

Angiogenesis is the formation of new blood capillaries from pre-existing blood vessels (1,2). Inflammatory cytokine mediated responses and angiogenesis play an important role in many diseases, such as peripheral arterial disease (PAD), cancer, and ocular diseases, as well as regenerative medicine and tissue engineering. The essential role of blood vessels in delivering nutrients makes angiogenesis important in the survival of cells within tissues, including tumor growth. Targeting angiogenesis is an important strategy in many contexts, for example, promoting blood vessel formation has been studied extensively for tissue engineering to support the long-term viability of engineered tissue constructs (3). Also since PAD is a manifestation of atherosclerosis, which leads to an obstruction in arteries and further limits blood flow to distal tissues (4), promoting angiogenesis has been an important investigational strategy for PAD treatment; however, it has not been successful. At least one potential explanation for the inability to modulate angiogenesis is that this process triggers inflammatory responses (4,5) that can limit the angiogenic response. Specifically, endothelial cells in response to inflammatory cytokines, such as IL-6, get activated and lead to increased vascular leakage, leukocyte recruitment, and further an accumulation of plaques, which blocks blood flow (6,7). On the other hand, inflammation can promote angiogenesis in many ways as well. Specifically, inflammatory tissues are often hypoxic which induces angiogenesis (8). Also, cells involved in inflammatory processes such as macrophages and fibroblasts secrete angiogenic factors that promote vessel formation (8). In addition, there is evidence that pro-inflammatory cytokines such as interleukin-6 (IL-6) and tumor necrosis factor alpha (TNFα) promote angiogenesis (8–10). Thus, inflammation is often associated with angiogenesis (8) and it plays an important role in the development of many diseases, such as cancer and PAD. Thus, the goal of this study is to investigate, in mechanistic detail using a computational model, IL-6 mediated signaling in vascular endothelial cells.

Numerous experimental and computational studies that investigated the array of responses to inflammatory cytokines in different cell types such as macrophages (11,12), T cells (13,14), and cancer cells (15,16). Also, recent reviews focused on computational models and analysis of angiogenic signaling (17,18). However, there is limited quantitative analysis of inflammatory together with angiogenic responses in endothelial cells to inform potential treatments that target inflammation and angiogenesis. Therefore, we aim to focus on inflammatory signaling in endothelial cells to characterize endothelial inflammatory and angiogenic responses. Many cytokines, such as IL-6, TNFα, and interleukin-1β (IL-1β) regulate inflammatory signaling (19–23). The role of many circulating biomarkers, such as selectins and interleukins in PAD has been reviewed (6,24). Also, potential anti-inflammatory strategies are reviewed for cardiovascular disease (25,26). In this study, we will focus on the intracellular signaling mediated by one of the major inflammatory cytokines, IL-6, as it has been identified as an important biomarker in inflammation in many diseases such as cardiovascular disease including PAD and cancer (6,24,25,27). In addition, elevated levels of IL-6 (28–33) and soluble IL-6 receptors (sIL-6R) (33,34) have been demonstrated in pathological conditions, including PAD and cancer.

Interestingly, IL-6 can act as both pro- and anti-inflammatory factor (27). IL-6 signaling transduces via binding to its membrane bound receptor (IL-6R) is referred to as classic signaling. When IL6 binds to its soluble receptor sIL-6R, and then recruiting glycoprotein 130 (gp130), this is referred to as trans-signaling (27). It has been shown that IL-6 classic signaling is associated with anti-inflammatory and regenerative responses, while IL-6 trans-signaling is involved in pro-inflammatory responses (27,35). Specifically, IL-6 binds to its receptors (IL-6R and/or sIL-6R) and gp130 and initiates signaling through the signal transducer and activator of transcription 3 (STAT3), mitogen-activated protein kinase (MAPK) and phosphatidylinositol 3-kinase/protein kinase B (PI3K/Akt) pathways to phosphorylate STAT3, extracellular regulated kinase (ERK) and Akt, respectively. The phosphorylated STAT3 (pSTAT3) and Akt (pAkt) are important signaling species in the inflammatory responses (36), while pAkt is believed to play an important role in cell survival (37–41) and phosphorylated ERK (pERK) is critical in cell proliferation (42,43), which are important processes involved in angiogenesis. Thus, we mainly focus on IL-6 trans-signaling mediated pSTAT3 and pAkt responses as indicators for pro-inflammatory signaling, and IL-6 classic signaling mediated Akt and ERK activation as signaling species for pro-angiogenic responses.

Given the complexity of biochemical reactions comprising inflammatory signaling networks, a better understanding of the dynamics of these networks quantitatively is beneficial for current anti-inflammatory strategies targeting endothelial cells. Computational modeling serves as a powerful tool to investigate molecular responses systematically. For example, Maiti et al. constructed a computational model to characterize the interactions between TNFα and interleukin-10 (IL-10) to study the interplay between pro- and anti-inflammatory signaling in macrophages (11). Reeh et al. developed a mathematical model to investigate IL-6 trans- and classic signaling in human hepatoma cells on a molecular level (44). In addition, Sadreev et al. built a mathematical model to describe the multisite phosphorylation for inflammatory signaling in T cells to study the mechanism of STAT3 and Interferon Regulation factor 5 (IRF-5) signaling in T cell differentiation (45). Cheong et al. studied the nuclear factor kappa B (NF-κB) signaling as it is important in inflammation and immune activation using an in silico model (46). Furthermore, Zhao et al. developed a large-scale mechanistic model which focused on seven driving pathways including interferon gamma (IFNγ), IL-1β, IL-10, IL-4, TNFα, hypoxia, and VEGF to characterize macrophage polarization (47). Later, Zhao et al. constructed a multiscale model that considers inflammatory signaling and includes intracellular, cellular, and tissue-level features to study the dynamic reconstitution of perfusion during post hindlimb ischemia (48).

Therefore, we constructed a computational model to characterize the intracellular signaling mediated by IL-6 in endothelial cells. Our work is the first model that focuses on IL-6 mediated signaling in endothelial cells to characterize endothelial inflammatory and angiogenic responses. The model predicts the dynamics of pSTAT3, pAkt and pERK in response to IL-6 classic and trans-signaling. The model predicts that IL-6 trans-signaling induces stronger downstream signaling and promotes inflammatory responses. Also, IL-6 trans-signaling plays a dominate role in the overall effects. In addition, both IL-6 and sIL-6R levels regulate signaling strength. Using this model, we also identified the influential species and kinetic parameters that specifically modulate the pSTAT3, pAkt, and pERK responses, which represent potential targets for inflammation- and angiogenesis-based therapies, and investigated their efficacy. The model predictions provide mechanistic insight into IL-6 signaling in endothelial cells. More broadly, this model provides a framework to study the efficacy of inflammation- and angiogenesis-based therapies for endothelial cells.

## MATERIAL AND METHODS

### Model construction

We constructed a molecular-detailed biochemical reaction network including IL-6 and their membrane-bound and soluble receptors, IL-6R and sIL-6R, respectively (Figure 1). Signaling is induced by the IL-6 binding to their receptors and gp130, culminating with phosphorylation of STAT3, Akt, and ERK through the STAT3, PI3K/Akt, and MAPK pathways. The molecular interactions involved in the network are illustrated in Figure 1. We adapted the IL-6 induced STAT3 pathway from the model developed by Reeh et al. (44), and we expanded the model by including PI3K/Akt and MAPK pathways from Song and Finley’s model (49). It is noteworthy that although STAT3 has been shown to have two phosphorylation sites, Tyr705 and Ser727 (50), it has been shown that IL-6 induced tyrosine phosphorylation depends on JAKs, while the mechanism of serine phosphorylation is not clear (51). Thus, we only considered the singly phosphorylated STAT3 (pSTAT3) in our model. In addition, we consider that activated Akt and ERK include both singly and doubly phosphorylated forms of each species since they have been reported to get activated at two phosphorylation sites (52,53). The model can be improved when more data are available. For simplicity, we collectively refer to these species as phosphorylated STAT3, Akt, and ERK (pSTAT3, pAkt, and pERK), respectively. The model reactions, initial conditions, and parameter values are provided in Supplementary Tables S1-3.

**Figure 1.**
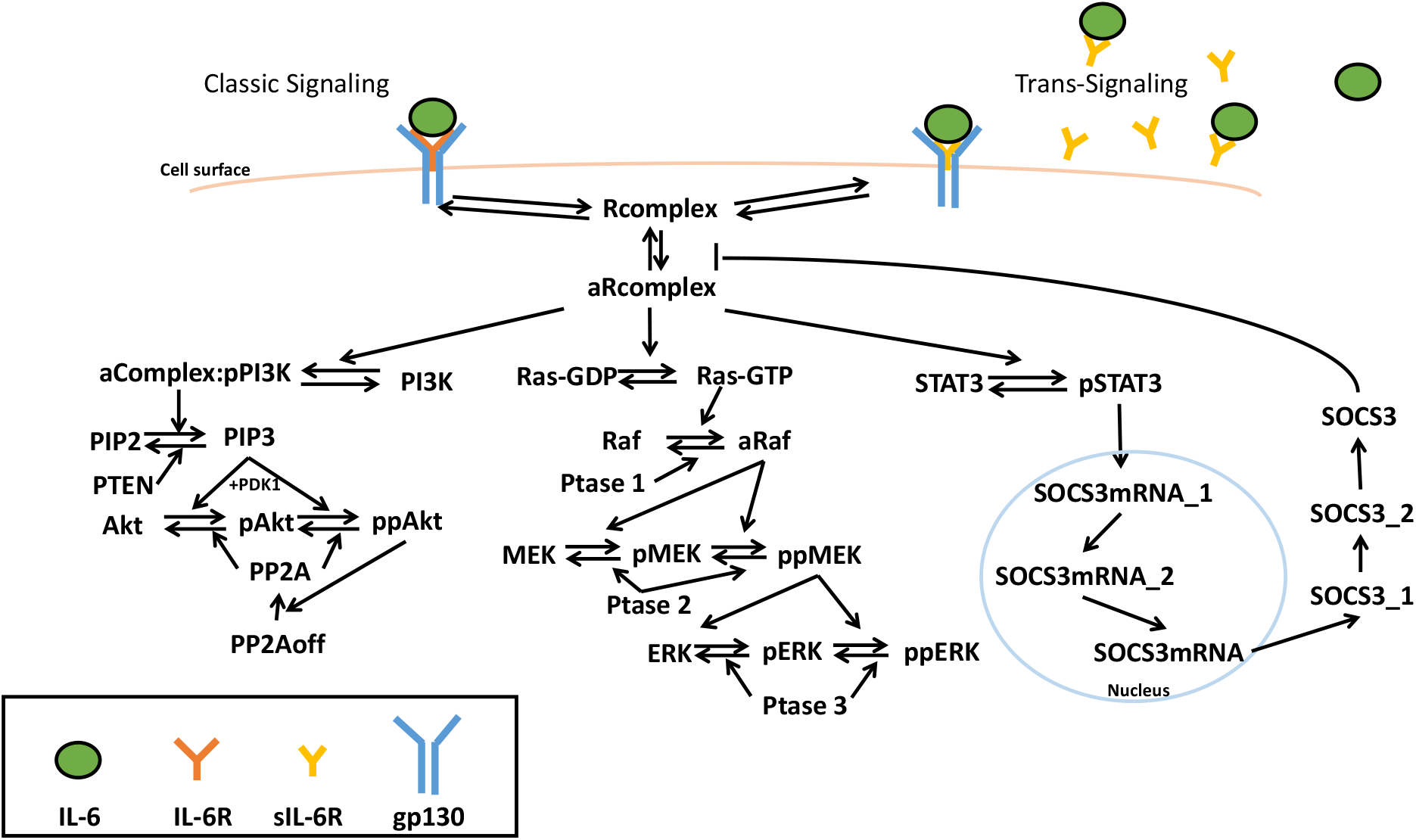
Schematic of IL-6 signaling network. IL-6 classic and trans-signaling is induced by IL-6 binding to membrane-bound and soluble IL-6 receptors, respectively, and recruiting gp130, which activates PI3K/Akt, MAPK, and STAT3 pathways and phosphorylates Akt, ERK, and STAT3, respectively.

The network is implemented as an ordinary differential equation (ODE) model using MATLAB (MathWorks, Natick, MA). The main model includes 55 reactions, 65 species, and 68 parameters. The initial variable settings of the initial conditions and parameters involved in IL-6 induced STAT3 pathway, and the variables involved in IL-6 induced Akt and ERK pathways are taken from the median values from Reeh et al.’s calibrated model (44) and Song and Finley’s fitted model (49), respectively. The reactions, initial conditions, and parameter values are listed in Tables S1 to S3. We listed four representative reactions below that describe the ligand-receptor binding as an example.

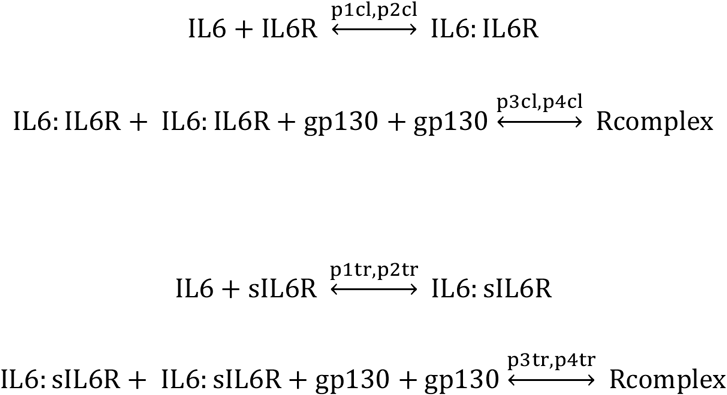

Because the simulated time is within four hours, we do not consider the degradation of the ligands or signaling species. The complete model is available in Supplemental File 3.

To set the initial conditions, since the expression of IL-6R in human endothelial cells is unclear (54), we corelated the IL-6R level with the gp130 level, which were measured in human umbilical vein endothelial cell (HUVEC) lysates (36) by one factor: ratio4 (*gp130*/*IL-6R* = 0.04 nM /0.0015 nM = 26) (Table S2). Also, we assumed a negligible basal sIL-6R in the system since the basal sIL-6R level (0.00019 nM) measured in HUVEC medium is much lower than IL-6R and gp130 measured in the HUVEC lysates (36), specifically the basal sIL-6R level is approximately 7.9-fold lower than IL-6R (0.0015 nM) and 210-fold lower than gp130 (0.04 nM).

### Sensitivity Analysis

To identify the parameters and initial concentrations that significantly influence the model outputs, we performed the sensitivity analysis to calculate the Partial Rank Correlation Coefficients (PRCCs), which indicate the correlation between the model inputs and model outputs (55). All targeted parameters and initial values were sampled simultaneously within specified bounds using Latin Hypercube Sampling (LHS); PRCC values for all targeted parameters and initial values were computed to evaluate the correlation between the model inputs (kinetic parameters or initial conditions) and the pSTAT3, pAkt, and pERK concentrations. In addition, the p-values from a t-distribution test corrected with Bonferroni correction were calculated. The PRCC values of the sensitive variables that are statistically significant (p-value < 0.05) were compared. The PRCC values can range from −1 to 1, where a higher positive PRCC value and a lower negative PRCC value indicate the input is more positively and negatively correlated to the output, respectively.

Before model training, we first calculated PRCC values for all the parameters and initial values. Since the parameters for STAT3 activation were adapted from Reeh et al.’s model (44), these variables were sampled using LHS within the estimated lower and upper bounds from Reeh et al.’s calibrated model (44) listed in Table S3. All remaining model parameters and initial values were sampled within two orders of magnitude above and two orders of magnitude below the baseline values, where the baseline values were taken from the median values estimated from published literature (49,56) listed in Tables S2-3. Based on the experimental data that were used for model training, we calculated the PRCC values for all the same concentrations and time points as those used in the experiments. The highest PRCC value (*PRCC*_*max*_) across all of the concentrations and time points was selected to represent the sensitivity index for each variable.

We also performed sensitivity analysis for the calibrated and validated model to identify potential targets for inflammation- and angiogenesis-based strategies.

### Identifiability analysis

In addition to parameter sensitivity, we also performed structural parameter identifiability analysis to consider the uncertainty caused by the model structure in a dynamical system to study molecular signal transduction. The identifiability analysis identifies the parameters that have one unique model output for each parameter value. In this methods, pair-wise correlation coefficients between parameters were calculated. The identifiable parameters have correlations with all other parameters between −0.9 and 0.9 while unidentifiable parameters have correlations of > 0.9 or < −0.9 with at least one other parameter.

### Data extraction

Data from published experimental study (36,57) were used for parameter fitting and model validation. Experimental data from plots were extracted using function *grabit*. The western blot data were extracted using ImageJ.

### Parameterization

A total of 35 influential variables with *PRCC*_*max*_ values greater than 0.4 and less than −0.4 were identified by sensitivity analysis. Of these, 28 identifiable variables were identified by identifiability analysis (Table S4, highlighted in red) Thus, we held the rest of the variables constant and estimated a total of 28 influential and identifiable variable values by fitting the model to experimental measurements (44) using Particle Swarm Optimization (PSO) implemented by Iadevaia et al. (58). We used MATLAB to implement the PSO algorithm. PSO starts with a population of initial particles (parameter sets). As the particles move around (i.e., as the algorithm explores the parameter space), an objective function is evaluated at each particle location. Particles communicate with one another to determine which has the lowest objective function value. The objective function for each parameter set was used to identify optimal parameter values. Specifically, we used PSO to minimize the weighted sum of squared residuals (WSSR):

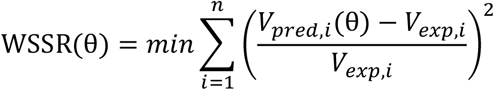

where V_*exp,i*_ is the *i*th experimental measurement, V_*pred,i*_ is the *i*th predicted value at the corresponding time point, and *n* is the total number of experimental data points. The minimization is subject to θ, the set of upper and lower bounds on each of the fitted parameters. The bounds for the parameters involved in the reactions for STAT3 activation were set to be the estimated lower and upper bounds from Reeh et al.’s calibrated model (44) and listed in Table S3. Also since the dissociation constant (Kd) of IL-6 for IL-6R has been reported to be 0.5 – 50 nM (44), we set the upper and lower bounds on p2cl to be 0.5*p1cl and 50*p1cl to confine the Kd for reaction IL-6 + IL-6R ⟷IL-6:IL-6R. In addition, the bounds for the remaining model parameters and initial values were set to be two orders of magnitude above and below the baseline parameter values, which were taken from the median values estimated from literature (49,56) and listed in Table S2-3.

The model was fitted using five experimental datasets from the literature (36), specifically: 1) relative change of pSTAT3 time course response from 0 to 240 min stimulated by 50 ng/ml IL-6 alone and in combination with 100 ng/ml sIL-6R compared with a reference point (pSTAT3 stimulated by 50 ng/ml IL-6 in combination with 100 ng/ml sIL-6R at 5 min); 2) relative change of ppAkt time course response from 0 to 240 min stimulated by 50 ng/ml IL-6 alone and in combination with 100 ng/ml sIL-6R compared with a reference point (ppAkt stimulated by 50 ng/ml IL-6 in combination with 100 ng/ml sIL-6R at 5 min); 3) relative change of pERK time course response from 0 to 240 min stimulated 50 ng/ml IL-6 alone and in combination with 100 ng/ml sIL-6R compared with a reference time point (pERK stimulated by 50 ng/ml IL-6 in combination with 100 ng/ml sIL-6R at 5 min); 4) relative change of pSTAT3 dose response stimulated by varying concentrations IL-6 from 0 to 50 ng/ml at 15 min compared with a reference point (pSTAT3 stimulated by 10 ng/ml IL-6 alone at 15 min); 5) relative change of pSTAT3 dose response stimulated by varying concentrations IL-6 from 0 to 50 ng/ml in combination with 100 ng/ml sIL-6R at 15 min compared with a reference point (pSTAT3 stimulated by 50 ng/ml IL-6 alone at 15 min). All published experiments were conducted using human umbilical vein endothelial cells (HUVECs) (36).

Model simulations were compared to experimental measurements. Specifically, the relative change of the responses was calculated as following:

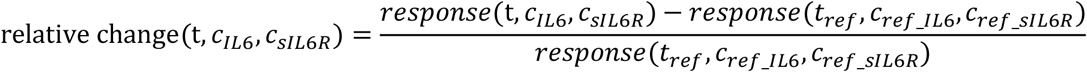

where *response*(*t, c*_*IL6*_, *c*_*sIL6R*_) is the level of pSTAT3, ppAkt, or pERK upon the stimulation of concentration *c*_*IL6*_ IL-6 in combination of concentration *c*_*sIL6R*_ sIL-6R at time t, and *response*(*t*_*ref*_, *c*_*ref*_*IL6*_, *c*_*ref*_*sIL6R*_) is the response (pSTAT3, ppAkt, or pERK) upon the stimulation of a reference concentration combination of *c*_*ref*_*IL6*_ IL-6 and *c*_*ref*_*sIL6R*_ sIL-6R at a reference time point *t*_*ref*_.

Here, the pSTAT3 in the model simulation includes all free and bound forms of singly-phosphorylated STAT3. Also, ppAkt includes all free and bound forms of doubly-phosphorylated Akt, since Zegeye et al. and Lindkvist et al. used anti-phospho-AKT^Ser473^ antibody for detecting phosphorylated Akt (36,57) and it has been reported that Akt gets phosphorylated at S473 as a secondary event (59–61). Thus, we compared the predicted doubly phosphorylated Akt (ppAkt) to experimental data (36,57). In addition, pERK in the model simulation includes all free and bound forms of singly- and doubly-phosphorylated ERK.

#### Constraints

In order to capture the whole dynamics of pSTAT3, pAkt, and pERK within 240 min, we applied a constraint for the relative change for ppAkt and pERK induced by IL-6 trans-signaling at 120 and 240 min by a factor of 0.1 when calculating the WSSR. Since the experimental relative change for ppAkt (0.32, and −0.0087) and pERK (0.14, and −0.26) induced by IL-6 trans-signaling at later time points are relatively low compared to other time points as they are reaching a plateau level after 100 min, we reduced their WSSR by a factor of 0.1 to let the model be more able to capture the whole dynamics rather than only the plateau behavior.

We first fitted the model 200 times to the experimental data. However, from the parameter sets that have the lowest errors, many fitted values were found at one of the bounds (Table S5). To exclude the possibility of arbitrary bounds limiting the parameter search space, we took the median values of 14 parameter sets that have the lowest errors as the baseline values and adjusted the bounds to be two orders of magnitude above and below the baseline parameter values (Table S6). The identified influential variables were estimated another 150 times with the new bounds. With the second round of fitting, none of the parameters were estimated to be at one of the bounds (Table S6). After model training, we validated the model with three datasets not used in the fitting. We predicted the 10 ng/ml IL-6 alone and in combination with 10 ng/ml sIL-6R induced pSTAT3, ppAkt, and pERK relative change time course responses using the reference points, pSTAT3, ppAkt, and pERK stimulated by 10 ng/ml IL-6 alone and in combination of 10 ng/ml sIL-6R at 10 min, respectively (57). The experiments (57) used for validation were performed using HUVECs.

#### Goodness of fit

The performance of the model was assessed as WSSR between the model predictions and experimental data and a runs test to determine if the predicted curve deviates systematically from the experimental data (62,63).

For all three datasets used for validation, we simulated the experimental conditions without any additional model fitting and compared to the experimental measurements. A total of 16 parameter sets with the smallest errors and p-values greater than 0.05 by performing the runs test were taken to be the “best” sets based on the model fitting and validation (Table S6) and were used for all model simulations. A p-value lower than 0.05 indicates the predicted curve deviates systematically from the experimental data, while a p-value greater than 0.05 suggests the residuals appear randomly distributed across the zero line (62,63).

### Monte Carlo simulations

To study the robustness of the system, the fitted model was run 1000 times by generating 1000 values for all parameters and non-zero initial concentrations, sampling from normal and lognormal distributions, respectively. For initial concentrations and parameters that were estimated by fitting to the experimental data, the mean values (μ) were the best fit, and for all other model variable values, we set μ to be the baseline values. The variances for the initial concentrations were set as an estimate of 10%μ. For all the parameters, we calculated the standard deviation (σ) to capture 99.7% of the possible values given the range of μ ± 50%μ (i.e., μ ± 3σ). It is worth noting that with this sampling, it is possible to get negative values, though this is unlikely to occur. However, if any negative values were selected, we resampled until all the sampled variables are positive.

### Signaling responses

We investigated the STAT3, Akt, and ERK phosphorylation responses upon stimulation by IL-6 classic- and/or trans-signaling.

#### Maximum pSTAT3, pAkt, and pERK

We calculate the maximum STAT3, Akt, and ERK phosphorylation levels induced by the stimulation by IL-6 classic- and/or trans-signaling within 4 hours.

#### Area under the curve (AUC) of pSTAT3, pAkt, and pERK

We calculate the AUC of STAT3, Akt, and ERK phosphorylation levels induced by the stimulation by IL-6 classic- and/or trans-signaling within 4 hours.

#### Reaction rates

We specify the rates of each reaction based on the law of mass action, where the rate of a chemical reaction is proportional to the amount of each reactant. For example, for the binding of IL-6 to IL-6R:

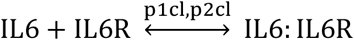

The reaction rate is

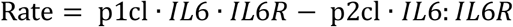

Here *p*1*cl* and *p*2*cl* are rate constants for the forward and reverse reactions, respextively, and *IL6, IL6R* and *IL6: IL6R* are the specie’s concentrations.

## RESULTS

### The fitted and validated molecular-detailed computational model captures the major characteristics of IL-6 induced STAT3, Akt, and ERK phosphorylation dynamics

For model training, we first identified the model variables (kinetic rates, initial concentrations, and factor ratio4) that significantly influence the model outputs, pSTAT3, pAkt, and pERK. To do so, we preformed sensitivity analysis using PRCC (see Methods for more details) and analyzed the PRCC values for all the species concentrations and kinetic rates. The highest PRCC values across all of the outputs and time points for a total of 65 species, 68 parameters, and 1 factor that affect initial concentrations, were compared, and 35 of them (Table S4 and Figure S1) were identified as influential to pSTAT3, pAkt, and pERK induced by 0 - 50 ng/ml IL-6 alone and/or with additional 100 ng/ml sIL-6R, which are the same concentrations applied experimentally (44). Of these, 28 of them were not correlated (highlighted in red, Table S4 and Figure S2), and we then estimated their values by fitting the model to experimental measurements (44) using PSO (58) (see Methods for more details).

The fitted model shows a good agreement with experimental results (Figure 2A-E). It quantitatively captures the dynamics of pSTAT3, ppAkt, and pERK by the stimulation of 50 ng/ml IL-6 alone (Figure 2A-C, light gray) and in combination with additional 100 ng/ml sIL-6R (Figure 2A-C, dark gray) (36). In addition, varying concentrations of IL-6 alone (Figure 2D) and in combination with additional 100 ng/ml sIL-6R (Figure 2E) induced-pSTAT3 dose responses have a good agreement with experimental measurements (36). The weighted errors for the 16 best fits range from 9.31 to 14.32 (Table S6).

**Figure 2.**
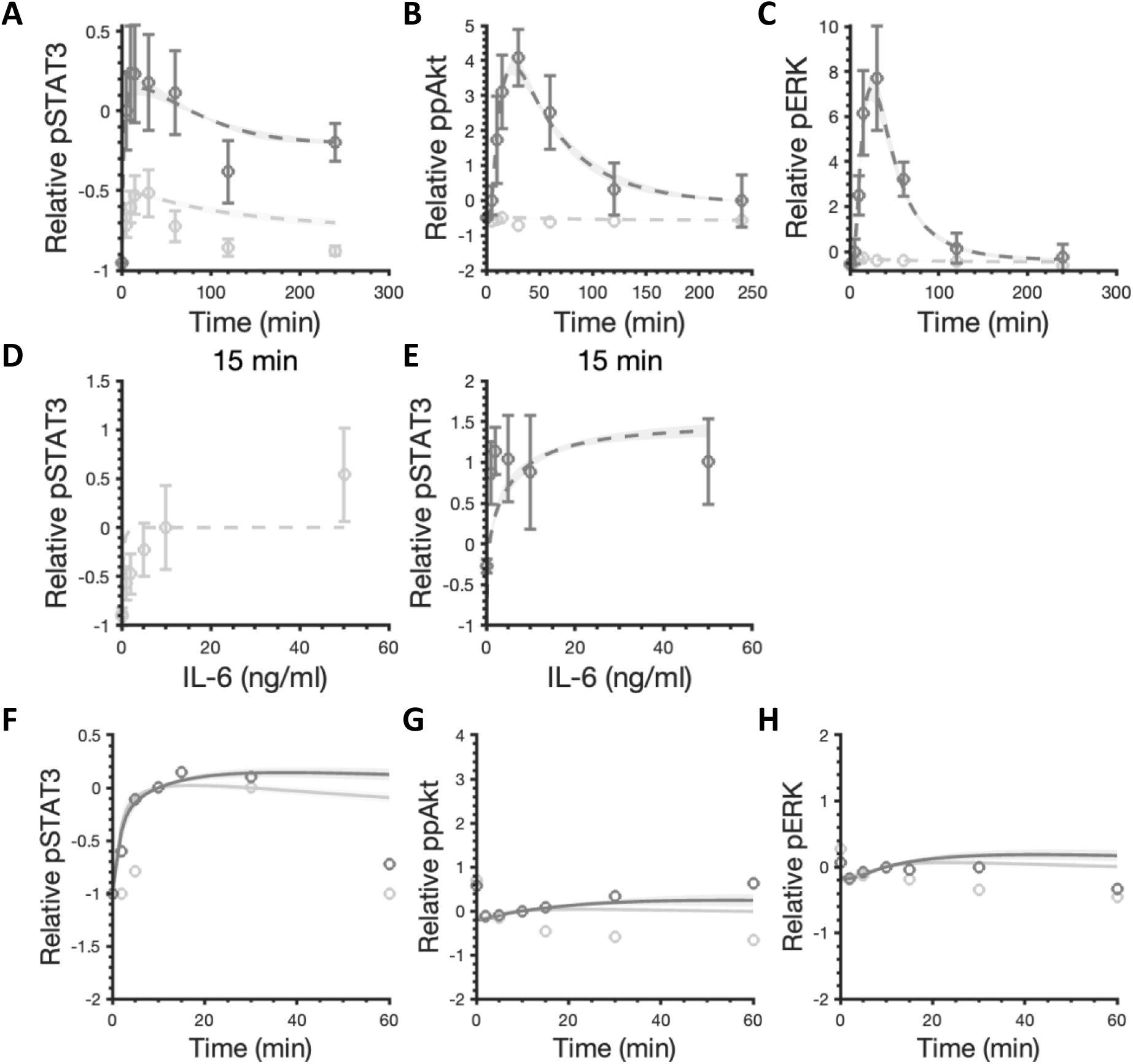
Model comparison to training and validation data for IL-6 stimulation. 50 ng/ml IL-6 with or without additional 100 ng/ml sIL-6R induced relative pSTAT3 (A), ppAkt (B), and pERK (C). Varying concentrations of IL-6 alone induced relative pSTAT3 (D) and with additional 100 ng/ml sIL-6R induced relative pSTAT3 (E). The circles are experimental data. Bars are mean ± SEM. Curves are the mean values of the 16 best fits. Shaded regions show 95% confidence intervals of the fits. Dashed and solid curves are training and validation results, respectively. Light gray: 50 ng/ml IL-6 (A-C), 0 – 50 ng/ml IL-6 (D), and 10 ng/ml IL-6 (F-H) stimulation; Dark gray: 50 ng/ml IL-6 + 100 ng/ml sIL-6R (A-C), 0 – 50 ng/ml IL-6 + 100 ng/ml sIL-6R (E), and 10 ng/ml IL-6 + 10 ng/ml sIL-6R (F-H) stimulation

In addition to model fitting, the model predictions are consistent with independent experimental observations that are not used in the model training (Figure 2F-H). To validate the model, we compared the model predictions to three independent sets of experimental data (57). Specifically, STAT3, Akt, and ERK phosphorylation by the stimulation of 10 ng/ml IL-6 alone (Figure 2F-H, light gray) and in combination with 10 ng/ml sIL-6R (Figure 2F-H, dark gray) matched the additional experimental measurements (57).

.It is noteworthy that we compared the predicted doubly phosphorylated Akt (ppAkt) to experimental data for model fitting and validation (36,57) (see Methods for more details). However, since both Akt T308 and S473 phosphorylation have shown to play an important role in the downstream signaling (52), we considered both singly and doubly phosphorylated forms of Akt to study its activation in the remainder of this work.

We performed Monte Carlo simulations (see Methods for more details) to study the predicted pSTAT3, pAkt, and pERK levels given variability in the initial concentrations and parameters. The model predictions with parameters values randomly varied within the range of the estimated values can still capture pSTAT3, pAkt, and pERK dynamics stimulated by IL-6 alone and in combination with sIL-6R (Figure S3). These simulations suggest that the overall dynamics of the model outputs, pSTAT3, pAkt, and pERK, are relatively robust to variability or uncertainty in initial concentrations and parameters in the signaling network.

### IL-6 classic and trans-signaling induced responses are dose-dependent

We first applied the experimentally validated model to explore the effects of IL-6 classic and trans-signaling on STAT3, Akt, and ERK phosphorylation. We found that the maximum pSTAT3, pAkt, and pERK levels within four hours increase with the increase of IL-6 concentrations (Figure 3A-C). IL-6-induced pSTAT3, pAkt, and pERK exhibit optimal ligand levels for inducing maximum responses as their dose response plateaus approximately at 0.2 nM by the stimulation of ligand concentration in the range of 0 nM – 1 nM (Figure 3A-C). In addition, the area under the curve (AUC) is quantified for pSTAT3, pAkt, and pERK dynamics within four hours as well and they exhibit a similar dose-dependent behavior (Figure S4A-C).

**Figure 3.**
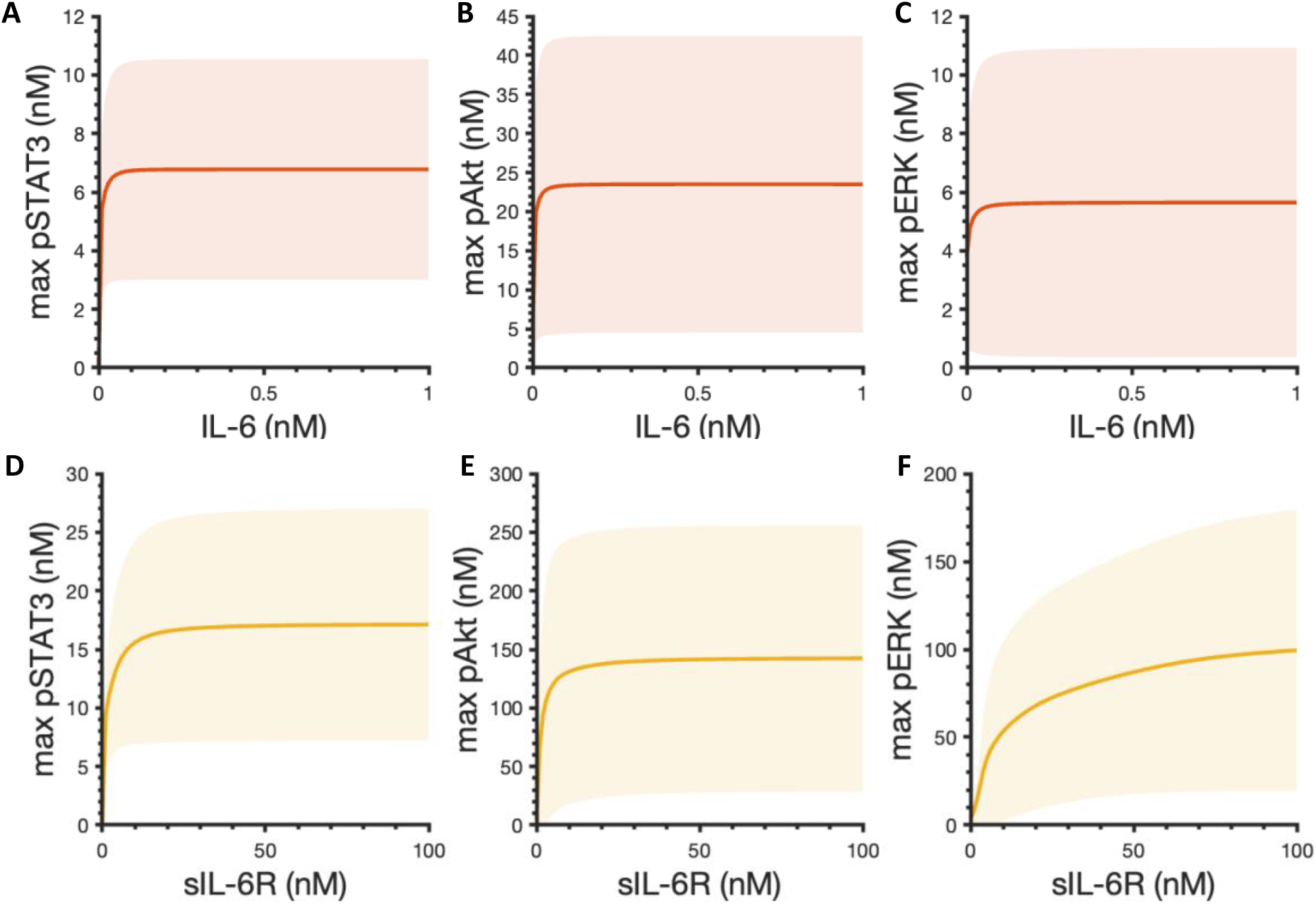
Predicted maximum pSTAT3, pAkt, and pERK responses. Maximum pSTAT3 (A), pAkt (B), and pERK (C) in response to IL-6 concentrations varying from 0 to 1 nM without sIL-6R. In the absence of IL-6R, 0.2 nM IL-6 in combination with sIL-6R concentrations varying from 0 to 100 nM induced maximum pSTAT3 (D), pAkt (E), and pERK (F). Curves are the mean values of the 16 best fits. Shaded regions show 95% confidence intervals of the fits. Orange: classic signaling responses; Yellow: trans-signaling responses.

We then set IL-6R level to be zero and simulated the phosphorylation of STAT3, Akt, and ERK in response to the stimulation of 0.2 nM IL-6 in combination with varying concentrations of sIL-6R to study the effects of IL-6 trans-signaling. A dose-dependent manner of STAT3, Akt, and ERK activation is also observed when considering the maximal phosphorylation levels (Figure 3D-F) and AUC (Figure S4D-F), respectively. Specifically, the maximum pSTAT3, pAkt, and pERK levels and AUCs increase with the increase of sIL-6R concentrations and show a trend of plateauing within 100 nM sIL-6R in combination with 0.2 nM IL-6 (Figure 3D-F and Figure S4D-F). In addition, since the maximum levels and AUCs exhibit the same trends as we observed (Figure 3 and Figure S4), for simplification, the maximum pSTAT3, pAkt, and pERK levels within four hours are utilized as indicators for pSTAT3, pERK and pAkt responses in this study.

### IL-6 trans-signaling induces stronger downstream responses compared to classic signaling and plays a dominant role in the overall effects

To compare the effects of classic and trans-signaling on STAT3, Akt, and ERK phosphorylation, we next set the concentration of sIL-6R at the same level as IL-6R for each fit, which is 28.84 nM on average among the 16 best fits, and simulated the dynamics of pSTAT3, pAkt, and pERK upon the stimulation of 0.2 nM IL-6 alone in the presence of IL-6R (orange) and 0.2 nM IL-6 in combination with a mean value of 28.84 nM sIL-6R in the absence of IL-6R (yellow) (Figure 4). Since IL-6R and sIL-6R are both present in the physiological and pathological conditions, we also studied the overall effects of the stimulation of 0.2 nM IL-6 in combination with 28.84 nM sIL-6R (mean) in the presence of IL-6R in pSTAT3, pAkt, and pERK responses (Figure 4, gray curves). We found that the IL-6 trans-signaling induced pSTAT3, pAkt, and pERK are significantly higher than the classic signaling induced responses, respectively (Figure 4). Also, IL-6 trans-signaling plays a dominant role in the overall effects in inducing pSTAT3, pAkt, and pERK as the overall effects induced responses overlap with the responses induced by the IL-6 trans-signaling (Figure 4).

**Figure 4.**
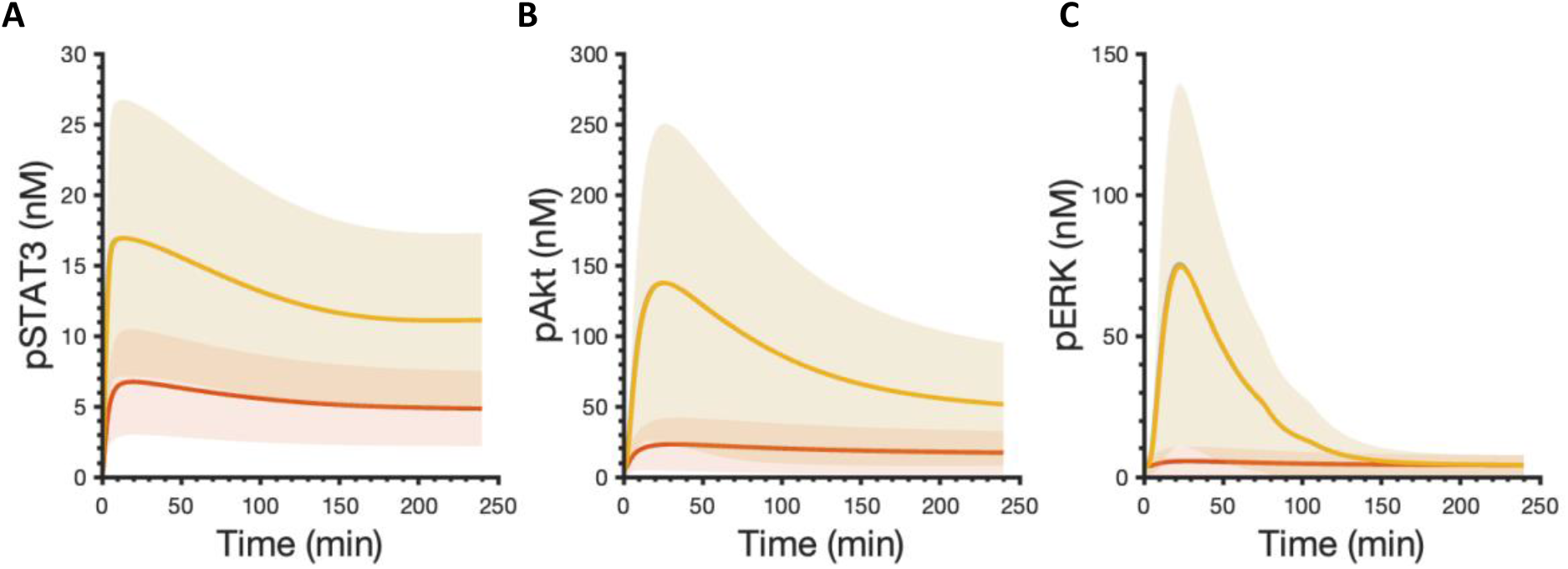
Predicted time courses of pSTAT3 (A), pAkt (B), and pERK (C) following stimulation by 0.2 nM IL-6 alone with a mean value of 28.84 nM IL-6R (orange), 0.2 nM IL-6 in combination with a mean value of 28.84 nM sIL-6R in the absence of IL-6R (yellow), and 0.2 nM IL-6 with a mean value of 28.84 nM of both IL-6R and sIL-6R (gray). Curves are the mean values of the 16 best fits. Shaded regions show 95% confidence intervals of the fits. Orange: classic signaling responses; Yellow: trans-signaling responses; Gray: overall responses.

To mechanistically explain this phenomenon, we explored the model structure and found that it is mainly caused by an assumption of a constant sIL-6R as our model input. Similar assumption was also made in Reeh’s model, specifically, Hyper-IL-6, which is a fusion protein composed of sIL-6R and IL-6 was applied to study the effects of IL-6 trans-signaling and it was assumed to be a constant model input as its concentration remained the same in the supernatant experimentally in an in vitro study (44). A sustained supply of soluble receptor leads to greater downstream responses compared to IL-6 classic signaling as the amount of IL-6R is limited. To verify this hypothesis, we set IL-6R as a constant input, which is the same as sIL-6R, in our model and compared the effects of classic and trans-signaling. We found that pSTAT3, pAkt, and Perk induced by IL-6 classic and trans-signaling almost overlap when IL-6R and sIL-6R are set at the same level and remain constant within four-hour simulation time (Figure S5A and Table 1), which confirms our hypothesis that stronger downstream responses induced by IL-6 trans-signaling are mainly caused by the sustained supply of sIL-6R.

**Table 1.** Predicted maximum pSTAT3, pAkt, and pERK responses for the baseline model, the modified model when IL-6R was set as a constant input, when kinetic rates governing R1 and R2 to be the same as the corresponding kinetic rates for R3 and R4, and when both IL-6R was set as a constant input and kinetic rates governing R1 and R2 to be the same as the corresponding kinetic rates for R3 and R4. The units in the table are nM. R1: IL-6 + IL-6R ⟷IL-6:IL-6R; R2: 2 IL-6:IL-6R + 2 gp130 ⟷Rcomplex; R3: IL-6 + sIL-6R ⟷IL-6:sIL-6R; R4: 2 IL-6:sIL-6R + 2 gp130 ⟷Rcomplex.

|  |  | Max pSTAT3 | Max pAkt | Max pERK |
| --- | --- | --- | --- | --- |
| Baseline | Classic | 6.74 | 23.10 | 5.58 |
|  | Trans | 16.91 | 137.39 | 74.57 |
|  | Overall | 16.93 | 137.55 | 75.34 |
| Receptor | Classic | 16.57 | 133.77 | 63.50 |
|  | Trans | 16.91 | 137.39 | 74.57 |
|  | Overall | 17.03 | 139.11 | 81.39 |
| Params | Classic | 6.74 | 23.07 | 5.59 |
|  | Trans | 16.91 | 137.39 | 74.57 |
|  | Overall | 16.92 | 137.50 | 75.19 |
| Params& Receptor | Classic | 16.91 | 137.39 | 74.57 |
|  | Trans | 16.91 | 137.39 | 74.57 |
|  | Overall | 17.08 | 139.14 | 83.57 |

We also noticed some differences in the dissociation constant (Kd) for the ligand-receptor binding reactions induced by the IL-6 classic and trans-signaling. Specifically, the Kd for reaction 1 (R1: IL-6 + IL-6R ⟷IL-6:IL-6R; mean Kd = 0.53 nM) is lower than the Kd for reaction 3 (R3: IL-6 + sIL-6R ⟷IL-6:sIL-6R; Kd = 17.94 nM); while the Kd for reaction 2 (R2: 2 IL-6:IL-6R + 2 gp130 ⟷Rcomplex; Kd = 0.05 nM) is higher compared to the Kd for reaction 4 (R4: 2 IL-6:sIL-6R + 2 gp130 ⟷Rcomplex; Kd = 0.02 nM) (Figure S6). It suggests a tighter binding of IL-6 to IL-6R than sIL-6R, while IL-6:sIL-6R binds tighter to gp130 than IL-6:IL-6R. However, no obvious difference was observed in pSTAT3, pAkt, and pERK when we set the kinetic rates governing the ligand-receptor binding reactions for classic signaling (R1 and R2) to be the same as the corresponding kinetic rates for the ligand-receptor binding reactions for trans-signaling (R3 and R4) compared with baseline model predictions (Figure S5B, Figure 4, and Table 1). It indicates that although there are some differences in ligand-receptor binding reactions induced by the IL-6 classic and trans-signaling, specifically IL-6 binds tighter to sIL-6R and IL-6:sIL-6R binds tighter to gp130 compared to classic signaling, it shows no obvious effects in the downstream signaling: 0.1% decrease in max pAkt and 0.2% increase in max pERK compared to the baseline model predictions (Table 1).

Last, we set IL-6R and sIL-6R at the same level and they remain constant within four hours and kinetic rates governing R1 and R2 to be the same as the corresponding kinetic rates for R3 and R4 and predicted the dynamics of pSTAT3, pAkt, and pERK (Figure S5C). The activation of STAT3, Akt, and ERK induced by IL-6 classic was found to overlap with the corresponding responses induced by IL-6 trans-signaling.

Overall, the model suggests that IL-6 trans-signaling induces stronger responses than classic signaling, and it plays a dominant role in the overall effects. It is mainly due to the sustained supply of sIL-6R.

### sIL-6R enhances the downstream signaling and promotes inflammatory responses

We next compared reaction rates for reactions 1-4 with or without IL-6R and sIL-6R (Figure 5). We found that IL-6 binds to sIL-6R faster compared to IL-6R in the beginning as the reaction rate for R3 is higher than the reaction rate for R1 (Figure 5A and C). Also, IL-6:sIL-6R binds faster to gp130 than IL-6:sIL-6R as the reaction rate for R4 is higher than the reaction rate for R2 in the beginning (Figure 5B and D). In addition, the reaction rates for R3 and R4 are more sustained compared to R1 and R2 (Figure 5A-B and C-D) since there is a sustained supply of sIL-6. It is noteworthy that additional sIL-6R makes reactions rates for R1 and R2 become negative over time (Figure 5A-B and E-F), which suggests a faster dissociation of IL6:IL6R and Rcomplex compared to the association of IL-6, IL-6R, and gp130. It indicates that more IL-6 and gp130 are freed from binding to IL-6R and available for binding to sIL-6R and inducing trans-signaling. Also, reactions rates for R3 and R4 show no obvious difference when both IL-6R and sIL-6R present compared to the reactions rates for trans-signaling (Figure 5C-D and G-H). Together, it indicates that additional sIL-6R shifts the signaling towards trans-signaling, which promotes inflammatory responses and this is consistent with the dominant role of trans-signaling in the overall effects.

**Figure 5.**
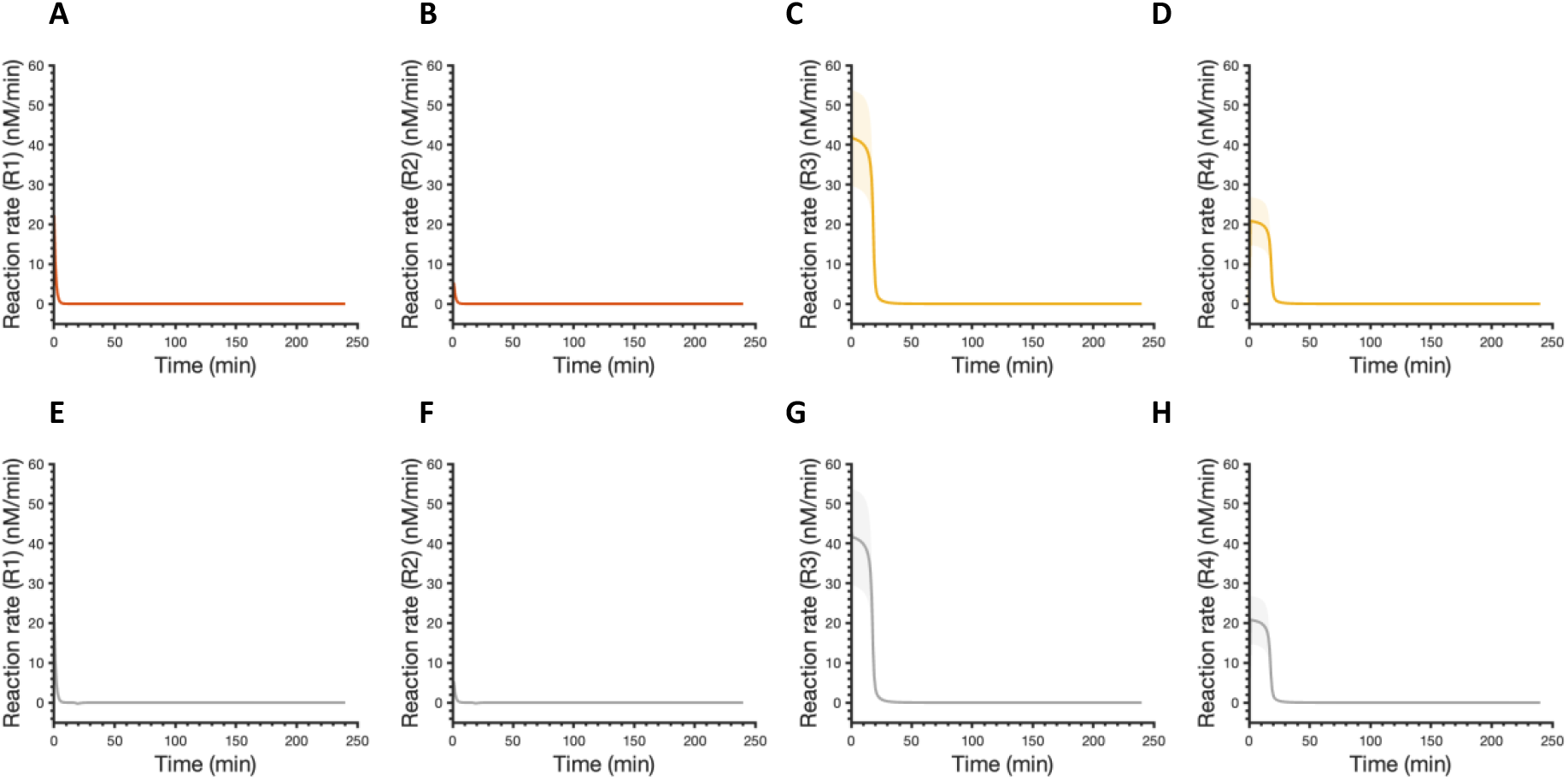
Reaction rates for ligand-receptor binding following stimulation by 0.2 nM IL-6 alone with a mean value of 28.84 nM IL-6R (orange) (A-B), 0.2 nM IL-6 in combination with a mean value of 28.84 nM sIL-6R in the absence of IL-6R (yellow) (C-D), and 0.2 nM IL-6 with a mean value of 28.84 nM of both IL-6R and sIL-6R (gray) (E-H). R1: IL-6 + IL-6R ⟷IL-6:IL-6R; R2: 2 IL-6:IL-6R + 2 gp130 ⟷Rcomplex; R3: IL-6 + sIL-6R ⟷IL-6:sIL-6R; R4: 2 IL-6:sIL-6R + 2 gp130 ⟷Rcomplex. Curves are the mean values of the 16 best fits. Shaded regions show 95% confidence intervals of the fits. Orange: classic

To further study the model details, we compared the time courses of relevant species involved in R1-R4 with or without IL-6R and sIL-6R (Figure S7). The model predicts that there is more IL-6:sIL-6R formed compared to IL-6:IL-6R (Figure S5A and C). Also, the predicted level of signaling Rcomplex induced by trans-signaling is higher than the classic signaling (Figure S5B and D). These model predictions further confirm that IL-6 trans-signaling induces stronger responses than classic signaling. Moreover, an accumulation of IL-6:IL-6R and a higher consumption of IL-6:sIL-6R induced by the overall effects compared to classic and trans-signaling respectively are observed (Figure S5A compared to E, and C compared to F). It is consistent with our predictions that additional sIL-6R shifts the signaling towards trans-signaling. In addition, the Rcomplex induced by the overall effects is approximately the same level as the Rcomplex induced by trans-signaling (Figure S5D compared to G), which agrees with the dominant role of trans-signaling in the overall effects.

Generally, the model suggests that IL-6 trans-signaling induces stronger responses and additional sIL-6R shifts the signaling towards trans-signaling, which promotes pro-inflammatory responses.

### Both IL-6 and sIL-6R levels regulate signaling strength

We next varied IL-6 and sIL-6R simultaneously and studied their combination effects in STAT3, Akt, and ERK activation. We found that there is a gradient towards the diagonal direction of increasing IL-6 and sIL-6R concentrations for each signaling species (Figure 6). As we observed previously, STAT3, Akt, and ERK activation plateau at approximately 0.2 nM IL-6 stimulation (Figure 3A-C), while additional sIL-6R further promotes the downstream signaling (Figure 6). Also, at a certain level of sIL-6R, adding IL-6 increases the STAT3, Akt, and ERK activation as well (Figure 6). An upregulation of IL-6 (31) and sIL-6R (64) has been reported in PAD conditions, which leads to stronger inflammatory responses. It is consistent with our model predictions as higher IL-6R and sIL-6R levels lead to greater phosphorylation of STAT3, Akt, and ERK (Figure 6).

**Figure 6.**
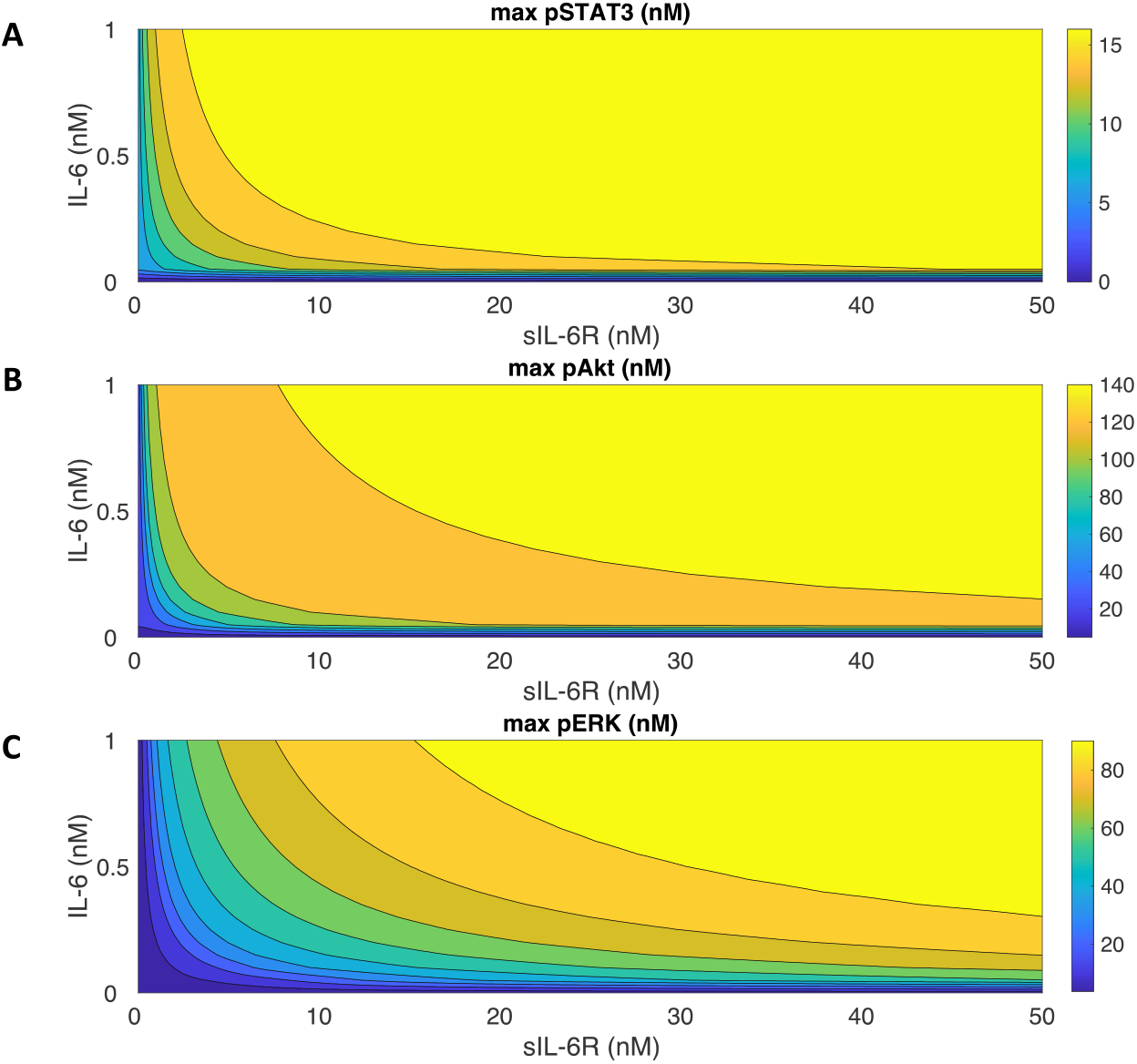
Predicted maximum pSTAT3, pAkt, and pERK responses with varying concentrations of IL-6 and sIL-6R. Maximum pSTAT3 (A), pAkt (B), and pERK (C) in response to the stimulation of 0 – 1 nM IL-6 in combination with 0 – 50 nM sIL-6R.

### Model identifies potential targets for anti-inflammatory and pro-angiogenic therapies and quantitively evaluates their efficacy

We performed sensitivity analysis using PRCC (see Methods for more details) for the experimentally validated model and identified influential initial concentrations (Figure S8A-C) and parameters (Figure S8D-E) to STAT3, Akt, and ERK activation. Specifically, all model parameters and initial values were sampled within two orders of magnitude above and two orders of magnitude below the baseline values. In this case, the baseline values for the fitted variables were the best fit estimated from model fitting. Based on the behaviors of max pSTAT3, pAkt, and pERK that reach a plateau as the IL-6 concentration increases (Figure 3), we selected 0.2 nM IL-6 as a representative concentration to capture the optimal responses induced by classic signaling.

Also, to compare the effects of IL-6 classic and trans-signaling, we took 32.58 nM sIL-6 as a representative concentration since it is the same level as the IL-6R concentration from the best fit. Therefore, we calculated the PRCC values for pSTAT3, pAkt, and pERK in response to the stimulation of 0.2 nM IL-6 in combination of 32.58 nM sIL-6R at eight time points (0, 5, 10, 15, 30, 60, 120, and 240 min) ranging from zero to 240 min. Again, the *PRCC*_*max*_ across all the concentrations and time points was compared for all the variables.

To analyze their effects in pSTAT3, pAkt, and pERK quantitatively, we varied each of identified influential variables within a finite range, specifically 10-fold above and below the baseline levels and compared with the baseline model predictions (Figure 7). When the ratio is greater than one, it suggests that varying the variable promotes the response; when the ratio is equal to one, it shows no effects on the response; when the ratio is less than one, it indicates an inhibitory effect on the response. We consider the effects of the perturbations as effective when the change of response is greater than 2-fold or less than 0.5-fold. We found that no initial concentration or parameter was observed to influence pSTAT3/STAT3 significantly (Figure 7A-B). In addition, Akt phosphorylation is positively regulated by PI3K and PIP2 levels (Figure 7C); while it is negatively regulated by STAT3, PTEN, and PP2A levels (Figure 7D). This is intuitive as PI3K and PIP2 are important signaling upstream species for Akt phosphorylation. PTEN and PP2A are phosphatases for PIP3 and pAkt. The impact of STAT3 level in the Akt phosphorylation is due to the competition between STAT3 signaling and Akt pathway. Also, parameter k_aAkt positively regulates pAkt, while k_aPTEN and k_aPP2A negatively regulate pAkt (Figure 7D) as k_aAkt is the association rate of PIP3 and Akt/pAkt, k_aPTEN is the association rate of PIP3 and PTEN association rate, and k_aPP2A is the association rate of pAkt/ppAkt and PP2A. Last, STAT3, PI3K, and Ptase2 negatively regulate ERK phosphorylation (Figure 7E). Because STAT3 and PI3K are signaling species involved in competitive pathways and negatively influence ERK activation. Ptase2 is the phosphatase for pMEK/ppMEK. Also, no parameter was observed to influence ERK phosphorylation significantly (Figure 7F).

**Figure 7.**
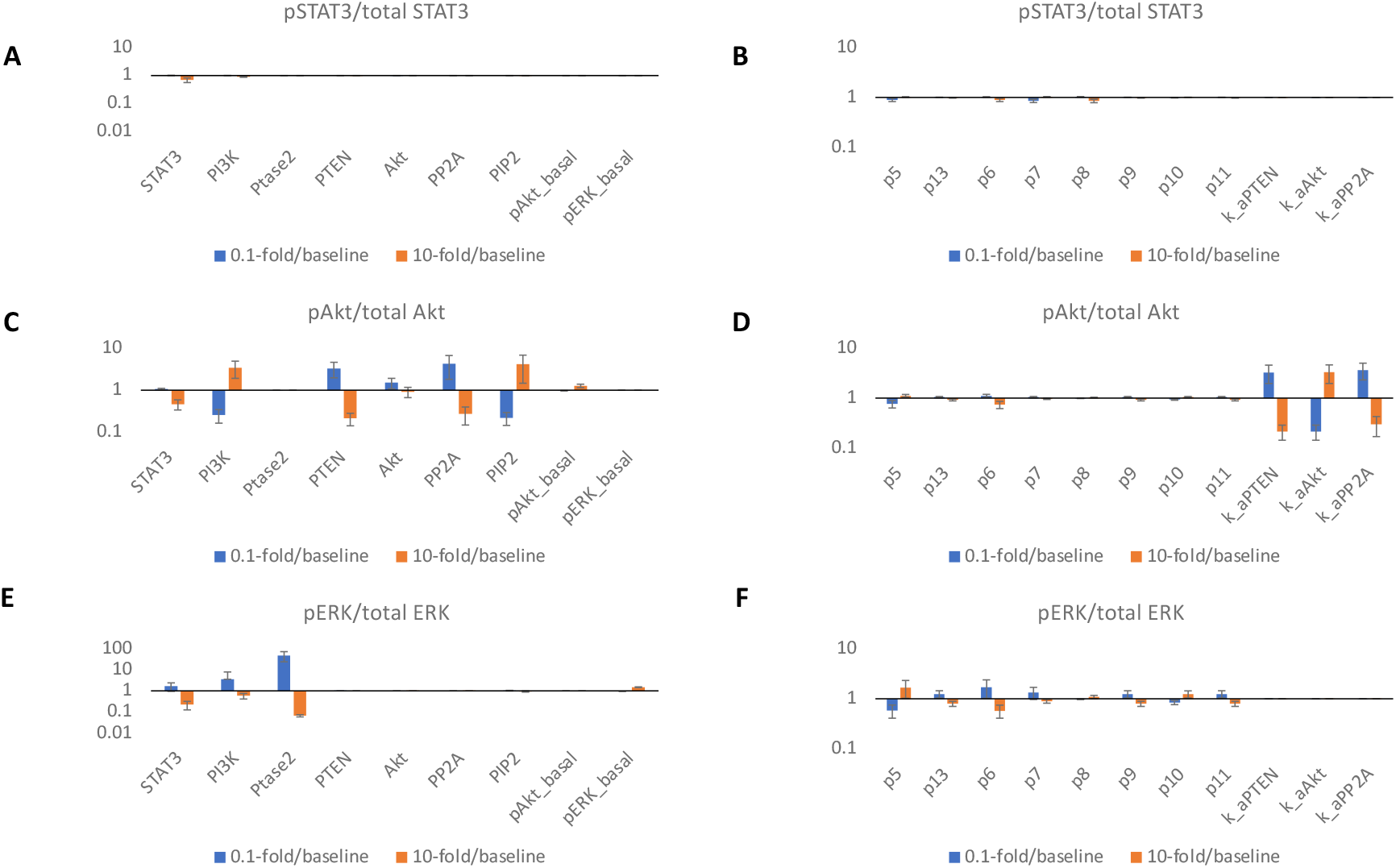
Predicted targets for modulating pSTAT3, pAkt, and pERK responses. *0*.*1-fold/baseline* (blue) and *10-fold/baseline* (orange) for 0.2 nM IL-6 with a mean value of 28.84 nM of IL-6R and 28.84 nM sIL-6R induced pSTAT3/total STAT3 (A, D), pAkt/total Akt (B, E), and pERK/total ERK (C, F) when varying identified influential initial concentrations (left) and parameters (right) by 0.1- and 10-fold of their baseline values. Bars are mean ± 95% confidence intervals of model predictions.

Thus, our model identifies potential targets for inflammation- and angiogenesis-based therapies and quantitively evaluated their efficacy.

## DISCUSSION

We developed an intracellular signaling model of IL-6 mediated inflammatory pathways in endothelial cells. The detailed computational model represents the reaction network of interactions on a molecular level. The model includes molecular interactions, kinetic parameters, and initial concentrations documented in literature, which are provided in the supplementary materials (Tables S1-3). Influential parameters were estimated by fitting the model to experimental data (36). Additionally, we validated the model using three independent experimental datasets (36).

IL-6 classic signaling is believed to be associated with anti-inflammatory or regenerative responses, while IL-6 trans-signaling is important in pro-inflammatory responses (35). It has been reported that IL-6 trans-signaling induces monocyte chemoattractant protein-1 (MCP-1) expression via activating STAT3 and Akt pathways but not MAPK signaling in HUVECs (36). Also, ERK activation is mainly believed to be important in cell proliferation (42). PI3K/Akt pathway has been reported to be critical in regulating cell survival and migration. Therefore, pSTAT3, pAkt, and pERK are main indices for inflammation and angiogenesis in this study. Specifically, this model focuses on IL-6 trans-signaling mediated pSTAT3 and pAkt responses as indicators for pro-inflammatory signaling, and IL-6 classic signaling mediated Akt and ERK activation as signaling species for pro-angiogenic responses.

The fitted model predicts pSTAT3, pAkt, and pERK responses upon the stimulation by IL-6 classic and/or trans-signaling. Overall, the model suggests that the max pSTAT3, pAkt, and pERK levels are IL-6 and sIL-6R dose-dependent. It has been shown that STAT3 phosphorylation in response to IL-6 classic and trans-signaling is dose-dependent in human hepatoma cells (HepG2) (44) and endothelial cells (36,54), which is consistent with our model predictions. In addition, our model predicts that IL-6 trans-signaling induces stronger responses and additional sIL-6R shifts the signaling towards trans-signaling and promotes inflammatory responses. It is consistent with other experimental work (36,54) and modelling work (44) that showed greater inflammatory response induced by IL-6 trans-signaling compared to classic signaling.

Angiogenesis and inflammation play an important role in many diseases, such as cancer, ocular, and cardiovascular diseases, including PAD. Angiogenesis also triggers inflammatory responses (4), which leads to malfunction of endothelial cells. Specifically, endothelial cells in response to pro-inflammatory cytokines, such as IL-6, get activated resulting in increased vascular leakage and leukocyte recruitment (6,7). However, there is a limited quantitative analysis of inflammatory pathways together with angiogenic responses in endothelial cells to inform potential treatments that target inflammation and angiogenesis. There are a number of computational models that study IL-6-induced signaling in many other cell types including hepatoma cells (44), cardiac fibroblasts (66,67), macrophages (68,69), and cancer stem cells (70). For example, Zeigler et al. built a large-scale mathematical model to characterize ten pathways including IL-6 signaling in cardiac fibroblasts to predict main regulators of fibrosis (66). In addition, Soni et al. developed a computational model that described IL-6 mediated macrophage activation in leishmaniasis (69). Moreover, Nazari et al. constructed a mathematical model to study IL-6 mediated cancer stem cell driven tumor growth (70). However, there is a limited quantitative understanding of IL-6 signaling in endothelial cells. Our research is the first computational model that focuses on IL-6 mediated signaling in endothelial cells to examine endothelial cytokine-mediated inflammatory and angiogenic responses.

Also, there are models that study cellular responses without considering intracellular signaling. For example, Nazari et al. linked cellular responses, specifically the temporal changes in the cancer stem cells, progenitor cells, and terminally differentiated cells with the fractional occupancy of bound receptor per cell (70). Our model can be utilized in combination with these types of models to more accurately predict cellular behaviors as more downstream signaling species could be better indicators for cellular responses.

This model can be beneficial to study the efficiency of angiogenesis- and inflammation-based therapies. Our model can identify the important variables to the pSTAT3, pAkt and pERK levels induced by IL-6 signaling and predict how pSTAT3, pAkt and pERK levels change by varying those parameters, which can provide quantitative insights into investigating the efficiency of targeting particular variables as angiogenesis- and inflammation-based strategies.

We do acknowledge certain limitations in our model. We adapted Reeh et al.’s IL-6 induced STAT3 pathway model which assumed that the ligand-receptor complex (Rcomplex) formed from classic and trans-signaling are the same. It implied a regulation of IL-6R and sIL-6R since the Rcomplex can associate and form both types of the receptor. Because the majority of the sIL-6R is generated by the shedding of the membrane bound IL-6R, and IL-6R expression can be regulated by ligands such as IL-6 in many cell types (71,72), we applied Reeh et al.’s model structure to include the potential regulation of the IL-6R and sIL-6R. Additionally, we simplified many species and reactions before activating STAT3, MEK/ERK and PI3K/Akt pathways by the stimulation of IL-6 because our main focus is their interactions. Also, we excluded soluble gp130 (sgp130) although their binding with IL-6:sIL-6R plays a role in inflammation (73). Their contributions to the model can be incorporated in future studies. In addition, due to the scarcity of the quantitative data on kinetics rates and initial conditions of IL-6 induced STAT3, Akt, and ERK activation in endothelial cells, we used parameters that govern IL-6 induced STAT3 pathway in human hepatoma cells (44) and VEGF- and FGF-induced Akt and ERK pathways in endothelial cells (49) as our initial guess to tune the parameters for IL-6 induced endothelial signaling, although the model was calibrated and validated using HUVEC data (36). Moreover, we assumed that IL-6 and sIL-6 levels are constant over four hours as the nutrients in the cell culture media are still sufficient. Same assumption has been made by Reeh et al. in their IL-6 signaling model to study human hepatoma cells (44). Also, since the expression of IL-6R is human endothelial cells is unclear (54), we estimated the receptor number which also causes a large variation in the model predictions. It can be improved when additional data on the receptor expression are available.

In conclusion, we developed a computational model to characterize the pSTAT3, pERK, and pAkt dynamics by the stimulation of IL-6 in endothelial cells. The model quantitatively studies STAT3, ERK, and Akt phosphorylation in response to IL-6 and sIL-6R and provides mechanical insight into inflammatory and angiogenic signaling in endothelial cells. The understanding of the regulation of inflammatory and angiogenic signals on a molecular scale can better aid the development of inflammation- and angiogenesis-based strategies.

## Acknowledgements

We are grateful to members of the Popel, Mac Gabhann, and Annex research groups for critical discussions, and especially Yu Zhang, Rebeca Oliveira. This work is supported by the National Institutes of Health grants R01HL101200 (ASP, BHA), R01HL141325 and 148590 (BHA) and R01CA138264 (ASP).

## Model availability

The model has been constructed using MATLAB (version R2021b). All model reactions, initial conditions, and values of parameters are provided in Tables S1 to S3 in Supplementary File 2 and MATLAB.m file containing model code is available in the Supplementary File 3.

## Author Contributions

MS and ASP conceived and designed the study. MS implemented the model in MATLAB, performed the computer simulations, analyzed the data, and drafted the manuscript. MS, ASP, YW, and BHA edited the manuscript.

## Supplementary Files

**Table S1. List of model reactions**.

**Table S2. List of species with non-zero initial concentrations**.

**Table S3. List of model parameters**.

**Table S4. PRCC values**

**Table S5. Fitted initial concentrations and parameters**.

**Table S6. Fitted initial concentrations and parameters with adjusted bounds**.

